# Molecular consequences of SCA5 mutations in the spectrin-repeat domains of β-III-spectrin

**DOI:** 10.1101/2024.09.17.613313

**Authors:** Sarah A. Denha, Naomi R. DeLaet, Abeer W. Abukamil, Angelica N. Alexopoulos, Amanda R. Keller, Alexandra E. Atang, Adam W. Avery

## Abstract

Spinocerebellar ataxia type 5 (SCA5) mutations in the protein β-III-spectrin cluster to the N-terminal actin-binding domain (ABD) and the central spectrin-repeat domains (SRDs). We previously reported that a common molecular consequence of ABD-localized SCA5 mutations is increased actin binding. However, little is known about the molecular consequences of the SRD-localized mutations. It is known that the SRDs of β-spectrin proteins interact with α-spectrin to form an α/β-spectrin dimer. In addition, it is known that SRDs neighbouring the β-spectrin ABD enhance actin binding. Here, we tested the impact of the SRD-localized R480W and the E532_M544del mutations on the binding of β-III-spectrin to α-II-spectrin and actin. Using multiple experimental approaches, we show that both the R480W and E532_M544del mutants can bind α-II-spectrin. However, E532_M544del causes partial uncoupling of complementary SRDs in the α/β-spectrin dimer. Further, the R480W mutant forms large intracellular inclusions when co-expressed with α-II-spectrin in cells, supporting that R480W mutation grossly disrupts the α-II/β-III-spectrin physical complex. Moreover, actin-binding assays show that E532_M544del, but not R480W, increases β-III-spectrin actin binding. Altogether, these data support that SRD-localized mutations alter key interactions of β-III-spectrin with α-II-spectrin and actin.

## Introduction

Mutations in the *SPTBN2* gene encoding the protein β-III-spectrin are associated with multiple neurological disorders including spinocerebellar ataxia type 5 (SCA5) and spinocerebellar ataxia autosomal recessive type 14 (SCAR14) (also known as spectrin-associated autosomal recessive cerebellar ataxia type 1 (SPARCA1)). SCA5 is caused by autosomal dominant missense or in-frame deletion mutations (1–6). Three SCA5 mutations were initially described in large kindreds of different nationalities (American, French and German) (1). Within these families, ataxic symptoms began in the third or fourth decade of life, on average, and were associated with cerebellar degeneration (7–9). In contrast, SCAR14/SPARCA1 is a recessive disorder, typically caused by mutations that truncate the β-III-spectrin protein (10–12). SCAR14/SPARCA1 causes cerebellar atrophy, severe infantile onset of ataxia and global motor and cognitive impairments. Recent clinical studies have also reported infants or adolescents, carrying *de novo* heterozygous dominant *SPTBN2* missense mutations, presenting with severe symptoms that mimic SCAR14 (13–23). These cases have been referred to as infantile SCA5 or *SPTBN2*-associated nonprogressive cerebellar ataxia (NPCA) (21), and here we will refer to these cases as early onset SCA5. The association of these mutations with severe, early onset symptoms suggests that these mutations more strongly or differently disrupt spectrin function, compared to the late onset SCA5 mutations. However, our current understanding of the molecular mechanisms of SCA5 pathogenesis is limited.

β-III-spectrin is a cytoskeletal protein with prominent expression in the soma and dendrites of cerebellar Purkinje neurons (24). β-III-spectrin is comprised of an N-terminal actin-binding domain (ABD), a central region containing 17 spectrin-repeat domains (SRDs), and a C-terminal pleckstrin homology domain (PH). β-spectrin proteins are known to form an antiparallel dimer with α-spectrin (25). Two heterodimers then associate to form a heterotetramer, which can cross-link actin filaments through the terminal β-spectrin ABDs (26–28). In cultured neurons, β-III-spectrin, like other neuronal β-spectrin proteins, was shown to form a sub-plasma membrane cytoskeleton composed of spectrin tetramers and actin filaments (29). Prior studies on erythrocyte β-I-spectrin and the broadly expressed β-II-spectrin indicate that dimerization critically depends on a high affinity, electrostatic interaction between N-terminal SRDs 1-2 of β-spectrin and the C-terminal SRDs 20-21 of α-spectrin (30–33). Following this nucleating interaction, the dimer is further stabilized by weak interactions between complementary SRDs that laterally associate along the two spectrin molecules (25, 30, 32, 33). It is also known that SRDs proximal to the ABD can enhance β-spectrin actin binding. Specifically, it was shown that a β-II-spectrin protein fragment containing the ABD and one or more SRDs attached, has increased actin-binding affinity compared to the ABD by itself (34).

SCA5 mutations cluster within the N-terminal ABD and neighboring SRDs. We previously showed that many ABD-localized mutations, including L253P (German kindred), cause increased actin binding (35, 36). The magnitude of the increase in binding varies among the mutations, and we suggested a correlation between magnitude of actin-binding affinity and age of onset/severity of symptoms. In contrast to the ABD-localized mutations, very little is known about the molecular consequences of the SRD-localized mutations. However, the position of the SCA5 mutations within the N-terminal SRDs suggests that they interfere with heterodimerization or actin binding. Here, we explored the molecular consequences of two SRD-localized SCA5 mutations: R480W, located in SRD2, and E532_M544del, located in SRD3. The R480W missense mutation is a *de novo* mutation reported in four unrelated patients with early (infantile) onset of symptoms including global developmental delay that leads to ataxia and intellectual disability (13–15, 20). E532_M544del was clinically characterized in the American kindred that descends from the paternal grandparents of the United States President Abraham Lincoln (7). E532_M544del causes an in-frame deletion of 13 amino acids in SRD3 and is associated with late onset SCA5. We performed biochemical and cell-based experiments to assess the impact of the mutations on β-III-spectrin folded state, dimerization with α-II-spectrin, and actin binding. Our findings support that the mutations differently alter how β-III-spectrin physically associates with α-II-spectrin, and that the E532_M544del mutation additionally impacts actin binding.

## Results

### Impact of R480W and E532_M544del on β-III-spectrin folded state and conformation

Numerous SCA5-associated missense and in-frame deletion mutations localize to SRDs 2-6 of β-III-spectrin, Fig. 1A. However, little is known about the molecular consequences of these mutations. Here we investigated the molecular consequences of the SRD2-localized R480W and SRD3-localized E532_M544del mutations. To study the impact of these mutations *in vitro*, we purified β-III-spectrin proteins containing the N-terminal ABD and first seven SRDs (amino acids 1-1066), Fig. 1B. For tandem affinity purification, the β-III-spectrin proteins were fused to an N-terminal His-SUMO tag and a C-terminal FLAG tag. The N-terminal His-SUMO tag was removed using Ulp1 protease, and the resulting purified β-III-spectrin proteins run at the expected molecular weight (∼122 kDa) following SDS-PAGE, Fig. 1B. Both the N-terminal ABD and SRDs are known to adopt highly α-helical folds (37–39). X-ray crystallography and cryo-EM structural data for spectrin proteins showed that the SRD consists of three α-helices (termed A, B, and C), bundled into an anti-parallel coiled-coil structure. This coiled-coil structure confers an overall rod-like shape to the SRD (40–44). Consecutive SRDs are linked to each other by a continuous helix formed by helix C of the first SRD and helix A of the second. To experimentally assess the folded state of the purified β-III-spectrin proteins, circular dichroism (CD) spectroscopy was performed. The wild-type protein shows a pronounced α-helical absorption profile characterized by two negative peaks near 208 and 222 nm, Fig. 1C. This absorption profile is thus consistent with the known α-helical fold of both the ABD and SRDs. Both R480W and E532_M544del mutant β-III-spectrin proteins also exhibit α-helical absorption profiles, indicating that the mutant proteins are also rich in α-helical content.

**Figure 1.**
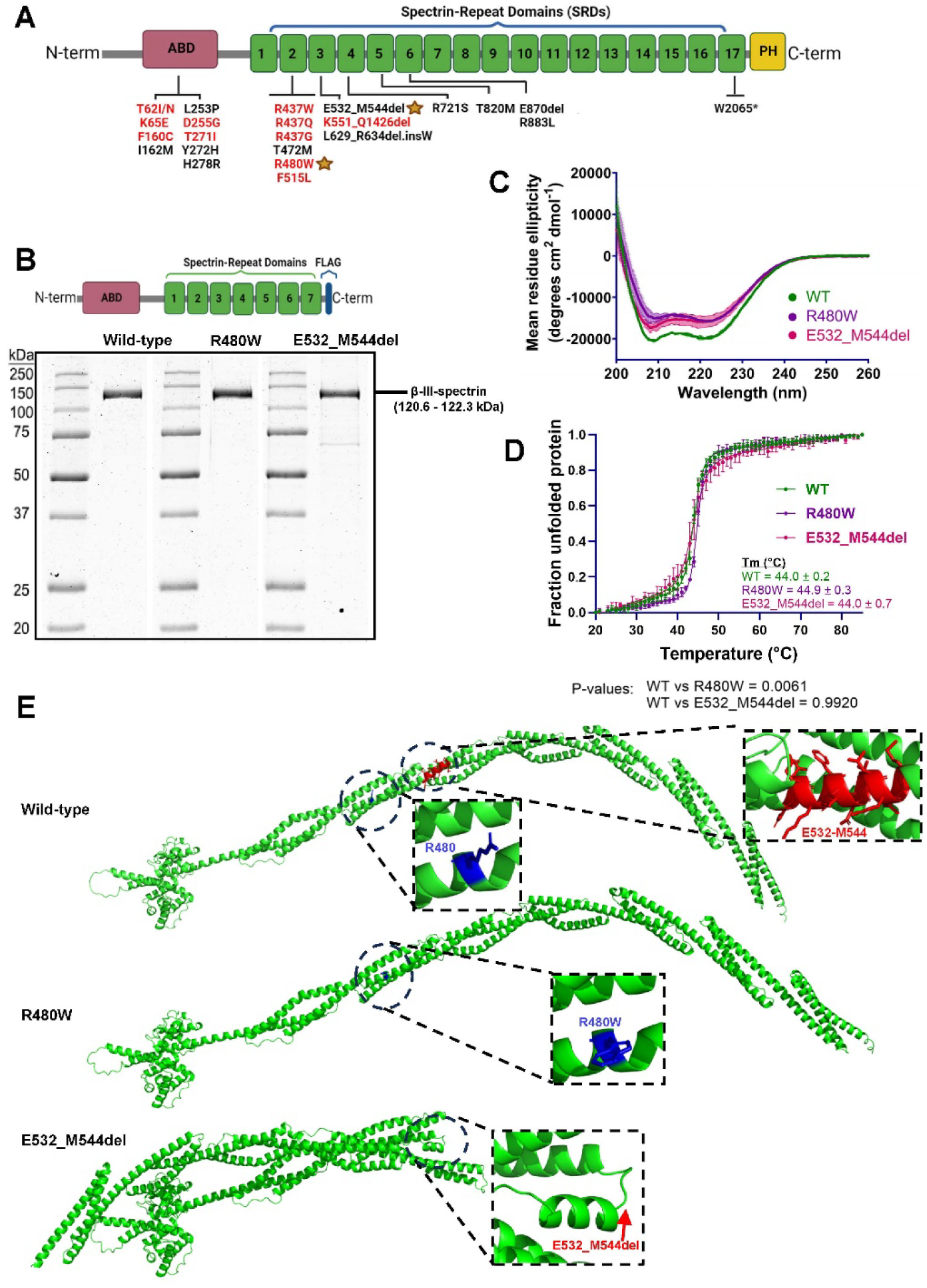
Structural analyses of R480W and E532_M544del β-III-spectrin proteins. (A) Domain diagram showing the N-terminal ABD, central 17 SRDs and C-terminal PH domain of β-III-spectrin. Many SCA5 mutations cluster in the ABD and the proximal SRDs (SRDs 2-6). Mutations in red are associated with early onset SCA5 and in black are associated with late onset SCA5. Y272H and W2065* were found together in a patient with early onset SCA5, as compound heterozygous variants Y272H/W2065* (18). SRD2-localized R480W and SRD3-localized E532_M544del mutations, which we characterized in this study, are marked with yellow stars. Domain diagrams were generated using BioRender.com (B) Top, domain diagram of the N-terminal fragment (amino acids 1-1066) of β-III-spectrin used in this study for *in vitro* biophysical and binding assays. The β-III-spectrin fragment is fused at the C-terminus to a FLAG epitope tag for purification purposes. Bottom, SDS-PAGE gel showing the purified β-III-spectrin proteins. The wild-type, R480W and E532_M544del mutant proteins run at the expected molecular weight (120.6-122.3 kDa). (C) CD α-helical absorption profiles measured over the range of 200-260 nm. The average of two different CD absorption profiles, using different protein preps, with error bars equal to standard deviation (D) CD melting curves measured at 222 nm. Each curve is the average of four independent denaturation curves, with error bars are equal to standard deviation. Tm values equal the average of four Tm values fitted in the replicate denaturation curves, with error given as standard deviation. Statistical significance was determined by Brown-Forsythe and Welch ANOVA tests followed by Dunnett’s T3 multiple comparisons test, n = 4. (E) AlphaFold structural predictions of wild-type and mutant β-III-spectrin proteins (1-1066 amino acids). R480 is located to helix B of SRD2 while E532-M544 residues are located at the start of SRD3 helix A. E532_M544del mutation is predicted to induce a disordered region to the continuous α-helix formed by SRD2 helix C and SRD3 helix A.

To assess how the mutations impact the stability of the β-III-spectrin proteins, thermal denaturation studies were performed while monitoring the CD peak absorption at 222 nm. Wild-type β-III-spectrin unfolded in a single sharp transition between two states, with a Tm of 44.0 ±0.2 °C, Fig. 1D. This melt curve is indicative of highly cooperative unfolding and a well-folded protein. Similarly, the melt curves for R480W and E532_M544del mutants show single step transitions with similar or slightly higher Tm values (44.9 ±0.3 °C for R480W and 44.0 ±0.7 °C for E532_M544del). The ∼1°C increase in Tm for R480W relative to wild-type was reproducible across multiple protein preparations, indicating that the R480W protein has a small stabilizing effect. The E532_M544del melt profiles repeatedly showed a less sharp/more gradual transition to the unfolded state, compared to wild-type and R480W mutant. This slight flattening of the melt curve suggests a modest reduction in cooperative unfolding (45) for the E532_M544del protein.

Complementing our CD structural studies, AlphaFold structure prediction software (46) was used to assess the impact of the R480W or E532_M544del mutation on the local SRD structure and overall protein conformation. In the wild-type protein, R480 localizes to helix B of SRD2. In contrast, the 13 amino acids deleted by E532_M544del mutation localize to the N-terminus of SRD3 helix A, Fig. 1E. The R480W mutant structure shows no loss in secondary structure relative to wild-type. Further, the predicted wild-type and R480W structures adopt similar overall conformations. In contrast, the E532_M544del mutation is predicted to replace the α-helical structure at the N-terminus of SRD3 helix A with an unstructured loop. The E532_M544del mutation thus eliminates the helical linker connecting SRD2 and SRD3. As a result, the unstructured loop introduces a hinge point, increasing conformational flexibility, such that the C-terminus of the protein can fold back towards the N-terminus. This loss of structured linker between SRD2 and SRD3 may account for partial reduction in cooperative unfolding of the E532_M544del mutant in our CD denaturation studies.

### R480 and E532-M544 residues are predicted to localize to the α/β dimer interface

Prior biochemical studies using N-terminal β-II-spectrin fragments and C-terminal α-II-spectrin fragments showed that β-II-spectrin SRDs 1-2 and α-II-spectrin SRDs 20-21 are necessary and sufficient for heterodimerization (33). A similar conclusion was drawn using N-terminal and C-terminal fragments of β-I-spectrin and α-I-spectrin, respectively (30,33). The high-affinity binding of β-spectrin SRDs 1-2 to α-spectrin SRDs 20-21 is thought to nucleate the pairing of additional β-spectrin and α-spectrin SRDs, strengthening the dimer. To explore whether the SRD-localized SCA5 mutations impact dimerization, we used AlphaFold to predict the structure of an α-II/β-III-spectrin dimer consisting of β-III-spectrin SRDs 1-4 and α-II-spectrin SRDs 18-21, Fig. 2A. In this structure, β-III-spectrin SRD2-localized R480 is closely positioned to α-II-spectrin SRD20 residues, Fig. 2B. A closer inspection shows that the R480 sidechain guanidinium group is predicted to form a charge-polar contact with the sidechain of α-II-spectrin residue N2109, and a salt bridge with α-II-spectrin E2108 sidechain, Fig. 2B. In addition, R480 is predicted to interact with both R437 and Y477 within β-III-spectrin SRD2. These contacts are supported by a recent cryo-EM structure of the blood α-I/β-I-spectrin dimer (44). In the α-I/β-I-spectrin dimer, the β-III-spectrin R480 equivalent, R477, interacts with α-I-spectrin N2059, equivalent of α-II-spectrin N2109, Fig. 2C. The equivalent residue to E2108 in α-I-spectrin (E2058) was not shown to interact with β-I-spectrin R477. However, the structure resolved similar interactions within β-I-spectrin, as R477 contacted R434 and Y474 (equivalent to β-III-spectrin R437 and Y477). Altogether, these analyses support that β-III-spectrin R480 participates in direct interactions with α-II-spectrin. This suggests that the R480W mutation is in position to alter binding of β-III-spectrin to α-II-spectrin. We similarly assessed the position of SRD3 amino acids E532-M544. In the AlphaFold structure, these 13 amino acids are positioned near the interface with α-II-spectrin SRD19, Fig. 2D. However, no direct interaction (< 4 Å) of these residues with α-II-spectrin is predicted. Moreover, the resolution of the equivalent region in α-I/β-I-spectrin dimer is poor, precluding a direct comparison in the experimentally derived structure. In sum, while E532-M544 residues are not predicted to directly contact α-II-spectrin, the position of the residues at the dimer interface supports that their deletion could disrupt dimer assembly.

**Figure 2.**
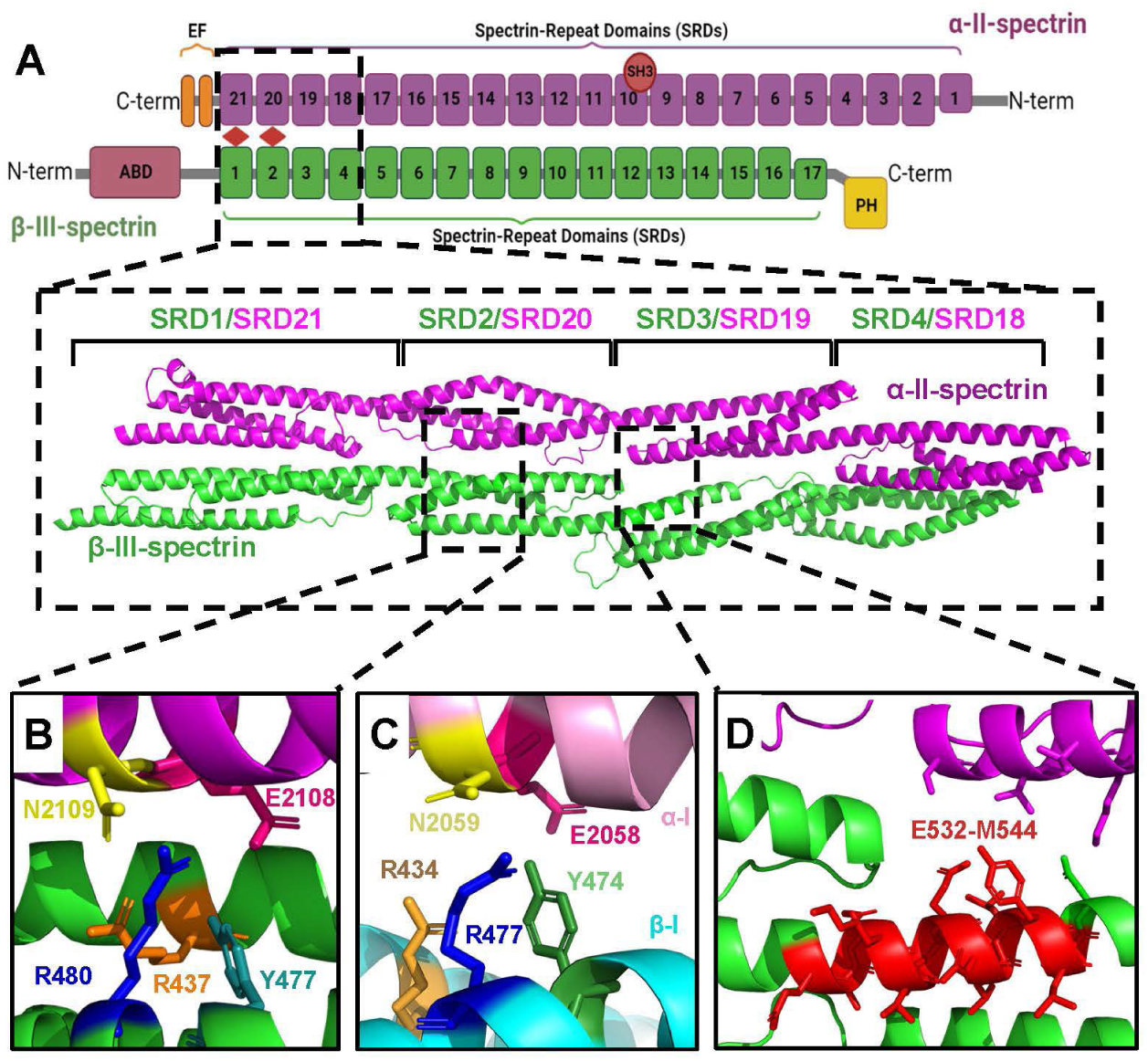
α/β-spectrin dimer structure. (A) Top, domain diagram of α-II/β-III-spectrin dimer. By analogy to other spectrin proteins, high affinity binding of N-terminal β-III-spectrin SRDs 1-2 to C-terminal α-II-spectrin SRDs 20-21, nucleate formation of an anti-parallel dimer in which complementary α-II/ β-III-spectrin SRDs laterally associate. Domain diagram generated using BioRender.com. Bottom, AlphaFold predicted structure showing the lateral association of SRDs 1-4 of β-III-spectrin with SRDs 18-21 of α-II-spectrin. Each SRD consists of a 3-helix bundle (helices A, B, and C). Consecutive SRDs are linked by a continuous helix formed by helix C of the first SRD and helix A of the next SRD. (B) Close-up view showing the location of R480 residue in SRD2 helix B and its predicted contacts in α-II-spectrin. R480 is predicted to interact with N2109 and E2108 residues on SRD20 of α-II-spectrin. Within β-III-spectrin, R480 is closely positioned to R437 and Y477. (C) Cryo-EM structure (PDB: 8IAH) of the erythrocyte α-I/β-I-spectrin dimer. The β-III-spectrin R480 equivalent in β-I-spectrin is R477.β-I-spectrin R477 is closely positioned to similar residues in β-I-spectrin and α-I-spectrin, supporting the accuracy of the AlphaFold model in panel B. (D) Close-up view showing the predicted location of E532-M544 residues at the N-terminus of SRD3 helix A of β-III-spectrin. E532-M544 residues are positioned at the interface with α-II-spectrin, although direct binding (< 4 Å) of these amino acids to α-II-spectrin is not predicted.

### E532_M544del partially opens α/β dimer interface

Several experimental approaches were taken to assess the impact of the R480W and E532_M544del mutations on binding of β-III-spectrin to α-II-spectrin. In one approach, we performed a cell-based FRET assay that provides a readout of dimerization. In this assay, HEK293T cells are transfected with a β-III-spectrin fragment (amino acids 1-1066), fused at the C-terminus to GFP (β-III-spectrin-GFP), together with the complementary fragment of α-II-spectrin (amino acids 1545-2473), fused at the N-terminus to mCherry (mCh-α-II-spectrin), Fig. 3A. Binding of these fragments is predicted to bring the C-terminal GFP of β-III-spectrin in close apposition to the N-terminal mCherry attached to α-II-spectrin. Indeed, cells co-expressing wild-type β-III-spectrin with α-II-spectrin generate an ∼8% FRET efficiency, Fig. 3A. In contrast, control cells expressing GFP (no β-III-spectrin fusion) together with mCherry-α-II-spectrin produced only a ∼2% FRET efficiency. Similar to wild-type, the R480W mutant produced an ∼8% FRET efficiency, suggesting that R480W does not impact formation of the dimer. In contrast, E532_M544del decreased FRET efficiency to a similar level as the GFP control, suggesting a strong decrease in dimerization.

**Figure 3.**
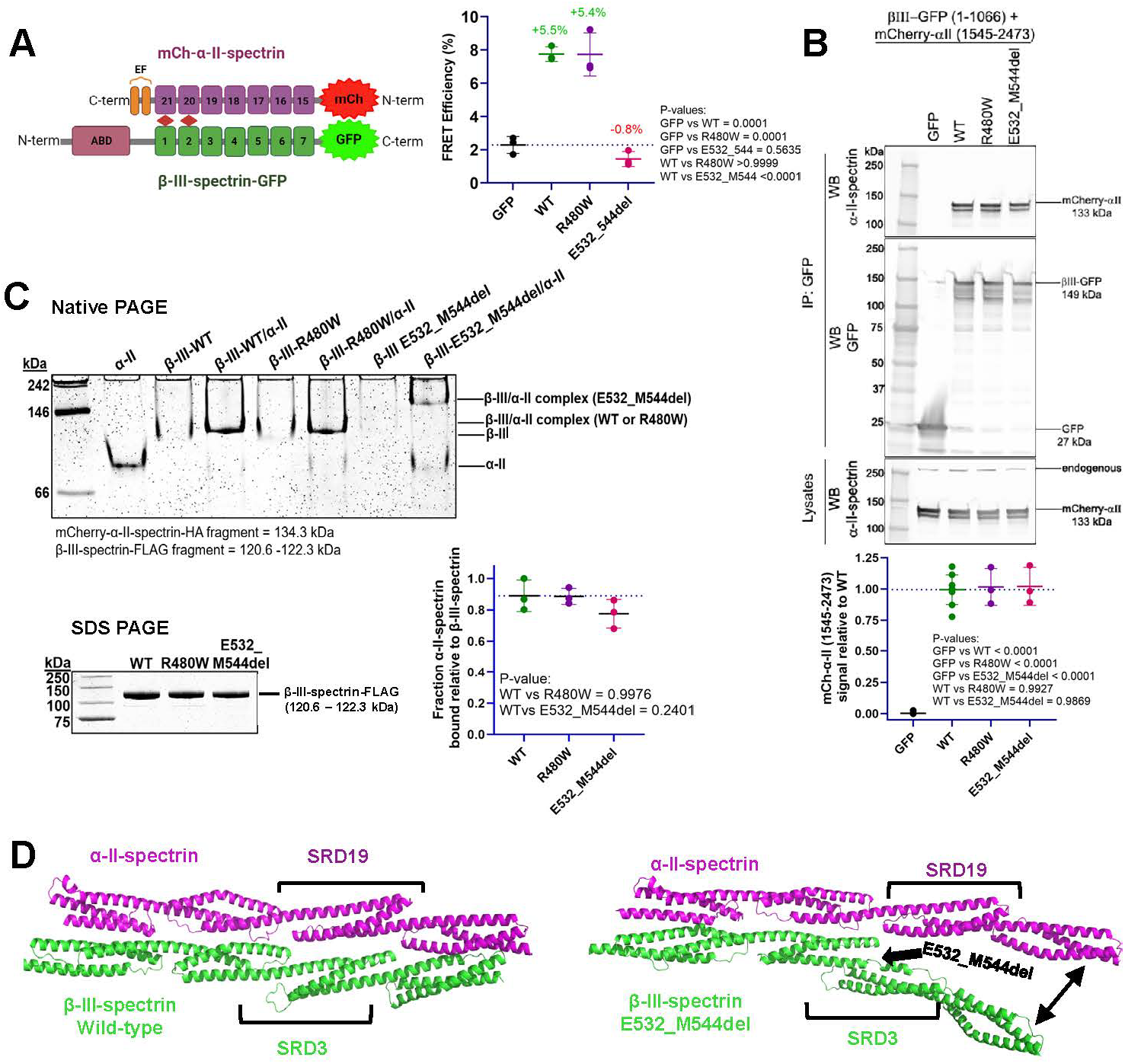
Cell-based, biochemical and *in silico* analyses of the impact of R480W and E532_M544del mutations on α-II-spectrin binding. (A) Live-cell FRET binding assays. Left, the FRET donor consists of β-III-spectrin fragment (amino acids 1-1066) fused at the C-terminus to GFP. The FRET acceptor contains α-II-spectrin fragment (amino acids 1545-2473) fused at the N-terminus to mCherry. Right, FRET binding data. (B) HEK293T cell co-immunoprecipitation (co-IP) assay using the β-III-spectrin and α-II-spectrin FRET assay proteins. Top, representative western blot of co-IP and lysate samples. Bottom, quantitation of α-II-spectrin protein in co-IP samples. (C) *In vitro* native PAGE mobility shift assay using purified, recombinant β-III-spectrin (amino acids 1-1066) and α-II-spectrin (amino acids 1545-2473) proteins. Top, representative native gel (Coomassie blue stained) showing the migration patterns of the α-II-spectrin C-terminal fragment in the absence of β-III-spectrin, β-III-spectrin N-terminal fragments in the absence of α-II-spectrin, and α-II/β-III-spectrin complexes. Protein concentration for β-III-spectrin or α-II-spectrin equaled to 0.5 µM. Bottom left, SDS-PAGE denaturing gel loaded with the individual β-III-spectrin proteins used in the native-PAGE binding assay. Bottom right, quantitation of depletion of the fast-running α-II-spectrin band in native PAGE binding assays, showing that both mutants bind a similar amount of α-II-spectrin compared to wild-type. (D) AlphaFold-Multimer predicted structures of the N-terminal SRDs 1-4 of β-III-spectrin bound to the complementary C-terminal SRDs 18-21 of α-II-spectrin for β-III-spectrin wild-type (left) versus the E532_M544del mutant (right). E532_M544del is predicted to open the interface between β-III-spectrin and α-II-spectrin beginning at the site of the mutation in SRD3. Statistical significance was determined using the Ordinary one-way ANOVA test followed by Tukey’s multiple comparisons test for FRET and co-IP assays and Dunnett’s multiple comparisons test for native PAGE assay, n = 3 independent experiments. Error bars equal standard deviation.

To confirm the decrease in dimer formation caused by E532_M544del, the co-association of the β-III-spectrin and α-II-spectrin proteins was assessed by co-immunoprecipitation (co-IP) binding assays. In the co-IP assays, the β-III-spectrin-GFP and mCh-α-II-spectrin proteins (same proteins used for FRET assays described above) were expressed in HEK293T cells. Wild-type and mutant β-III-spectrin proteins were immunoprecipitated using GFP antibody, and the co-associated α-II-spectrin protein levels were quantified using immunoblotting, Fig. 3B. Surprisingly, a similar amount of mCh-α-II-spectrin is present across the wild-type, R480W and E532_M544del mutant IP samples. This suggests that neither mutation causes a strong reduction in binding affinity of β-III-spectrin to α-II-spectrin.

We further tested binding of β-III-spectrin to α-II-spectrin using a biochemical, native PAGE mobility shift assay. Purified β-III-spectrin fragments (amino acids 1-1066), Fig. 1B, were incubated with the complementary α-II-spectrin fragment (amino acids 1545–2473). In native PAGE, control samples containing 0.5 µM α-II-spectrin or β-III-spectrin show that the α-II-spectrin protein migrates faster and in a more well-defined band compared to the β-III-spectrin proteins, Fig. 3C. Of the β-III-spectrin proteins, the E532_M544del mutant is most poorly resolved, supporting that the mutant adopts an altered conformation(s) compared to wild-type or R480W, in agreement with our AlphaFold prediction. In samples containing β-III-spectrin combined with α-II-spectrin, the α-II-spectrin and β-III-spectrin proteins shift up (migrate more slowly) to form a new single intense band, representing a physical complex between β-III-spectrin and α-II-spectrin. For wild-type and R480W, the α-II/β-III-spectrin complexes migrate at a similar position, located just above the β-III-spectrin band observed in the wild-type or R480W β-III-spectrin only control. In contrast, for E532_M544del, the α-II/β-III-spectrin complex migrates more slowly (at a higher position in the gel), suggesting a distinct conformation for the E532_M544del α-II/β-III-spectrin complex. Because the α-II-spectrin band was most well-defined and well-separated, the extent of β-III-spectrin binding to α-II-spectrin was determined by quantifying depletion of the α-II-spectrin band upon addition of β-III-spectrin. Notably, as a further loading control, the spectrin proteins were analysed by denaturing SDS-PAGE, Fig. 3C, bottom left. Quantitation of α-II-spectrin band depletion, relative to the amount of β-III-spectrin loaded, shows that the wild-type and mutant β-III-spectrin proteins bind a similar amount of α-II-spectrin. This *in vitro* assay is thus consistent with the similar binding of wild-type and mutant β-III-spectrin proteins to α-II-spectrin measured by our cell-based co-IP assays.

We further explored the interaction of the wild-type and E532_M544del β-III-spectrin (SRDs 1-4) with the α-II-spectrin (SRDs 18-21) using AlphaFold-Multimer (47). For wild-type, β-III-spectrin SRDs 1-4 lay alongside and antiparallel to α-II-spectrin SRDs 18-21, Fig. 3D. In contrast, for E532_M544del, only β-III-spectrin SRDs 1-2 are in contact with α-II-spectrin. Following the E532_M544del mutation, SRDs 3-4 are positioned away from α-II-spectrin. This suggests that the E532_M544del mutation does not disrupt nucleation mediated by binding of β-III-spectrin SRDs 1-2 to α-II-spectrin SRDs 20-21. However, the E532_M544del disrupts the coupling of β-III-spectrin SRDs to α-II-spectrin SRDs outside of the high affinity nucleation site. This structure prediction explains why E532_M544del mutation abolishes binding in our FRET assay that requires association of the β-III-spectrin C-terminal SRDs with α-II-spectrin, but does not impact overall binding in the co-IP and native PAGE assays. Further, the predicted open confirmation of the E532_M544del α-II/β-III-spectrin dimer explains the slow migration of the E532_M544del α-II/β-III-spectrin complex, relative to wild-type or R480W α-II/β-III-spectrin complex, in native PAGE.

### R480W and E532_M544del β-III-spectrin proteins co-localize with α-II-spectrin at the plasma membrane and cytoplasmic inclusions

Previously we showed in transfected HEK293T cells that full-length wild-type β-III-spectrin is predominantly localized to the plasma membrane (48). To determine whether the R480W or E532_M544del mutation impacts the subcellular localization of β-III-spectrin or α-II-spectrin, HEK293T cells were co-transfected with full-length, GFP-tagged β-III-spectrin proteins (GFP-β-III-spectrin) and full-length, mCherry-tagged α-II-spectrin (α-II-spectrin-mCherry). In agreement with our prior observation, wild-type β-III-spectrin shows prominent localization to the plasma membrane, Fig. 4. Further, α-II-spectrin co-localizes with wild-type β-III-spectrin at the plasma membrane (Pearson correlation coefficient ∼0.7). In contrast, in control cells expressing α-II-spectrin with a plasma membrane GFP tagged marker (PM-GFP), α-II-spectrin localization appears diffuse in the cytoplasm and less cortically localized (Pearson correlation coefficient ∼0.5). This suggests that β-III-spectrin recruits α-II-spectrin to the plasma membrane. The R480W and E532_M544del mutants also show pronounced co-localization with α-II-spectrin at the cell cortex (with Pearson correlation coefficients of ∼0.9), supporting that both mutants interact with α-II-spectrin for recruitment to the plasma membrane. Notably, for the R480W mutant, in addition to co-localizing with α-II-spectrin at the plasma membrane, large, spherical cytoplasmic inclusions are present that contain both the β-III-spectrin mutant and α-II-spectrin, Fig. 4A. Altogether, these data support that both full-length mutant proteins bind α-II-spectrin in cells.

**Figure 4.**
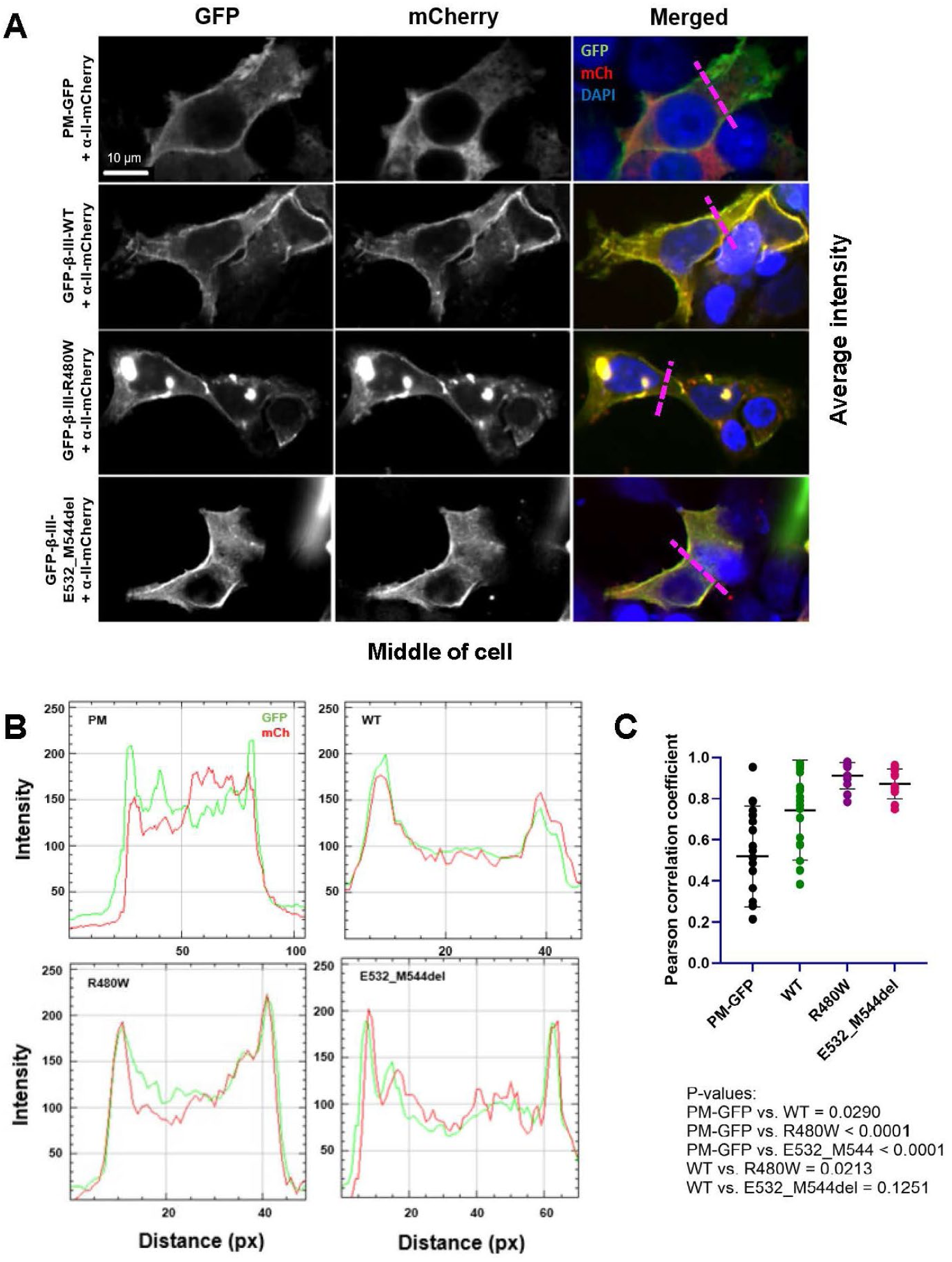
Wild-type and mutant β-III-spectrin proteins recruit α-II-spectrin to the plasma membrane. (A) Representative images of HEK293T cells co-expressing full-length GFP-tagged β-III-spectrin and mCherry-tagged α-II-spectrin. Each image is the average intensity of three z-stack planes in the middle of the cell. Wild-type β-III-spectrin localizes primarily to plasma membrane. α-II-spectrin shows pronounced co-localization with wild-type β-III-spectrin at the plasma membrane. In contrast, in cells expressing the plasma membrane-targeted GFP (PM-GFP), α-II-spectrin shows little plasma membrane localization and is instead diffusely localized in the cytoplasm. This suggests that wild-type β-III-spectrin recruits α-II-spectrin to the plasma membrane. Like wild-type, R480W and E532_M544del mutants also localize to plasma membrane and recruit α-II-spectrin to the plasma membrane. In addition to plasma membrane localization, R480W is also present in large cytoplasmic inclusions that contain α-II-spectrin. (B) Representative line scans of GFP and mCherry fluorescence intensities measured along the pink dash lines crossing the plasma membrane of the cells shown in A. Wild-type and mutant β-III-spectrin show peak intensities at the cell membrane which overlap with the α-II-spectrin peak intensities. In contrast, in PM-GFP control cells, α-II-spectrin intensities appear flatter along the measured distance. (C) Co-localization quantitation using Pearson correlation coefficient. GFP and mCherry fluorescence intensities were measured along lines crossing the plasma membrane, as shown in panel A. R480W and E532_M544del show similar or slightly higher Pearson correlation coefficients compared to wild-type. Statistical significance is determined using Brown-Forsythe and Welch ANOVA tests followed by Dunnett’s T3 multiple comparisons test, n= 11-25. Error bars equal standard deviation.

To determine if the R480W inclusions form in the absence of α-II-spectrin co-expression, the R480W mutant was expressed with and without α-II-spectrin in HEK293T cells. Interestingly, in cells not expressing α-II-spectrin, the R480W mutant shows little localization to inclusions, Fig. 5. The few R480W inclusions present appear small and irregularly shaped, similar to the occasional inclusions observed in cells expressing wild-type or the E532_M544del mutant (in the absence of α-II-spectrin). Indeed, in the absence of α-II-spectrin co-expression, the percentage of cells showing any β-III-spectrin inclusion was similar among wild-type, R480W and the E532_M544del mutant (∼50%), Fig. 5B. This supports that neither mutation causes internal accumulation of β-III-spectrin, which may result from possible protein destabilization/unfolding. This is consistent with our CD data showing that the mutations do not cause protein destabilization.

**Figure 5.**
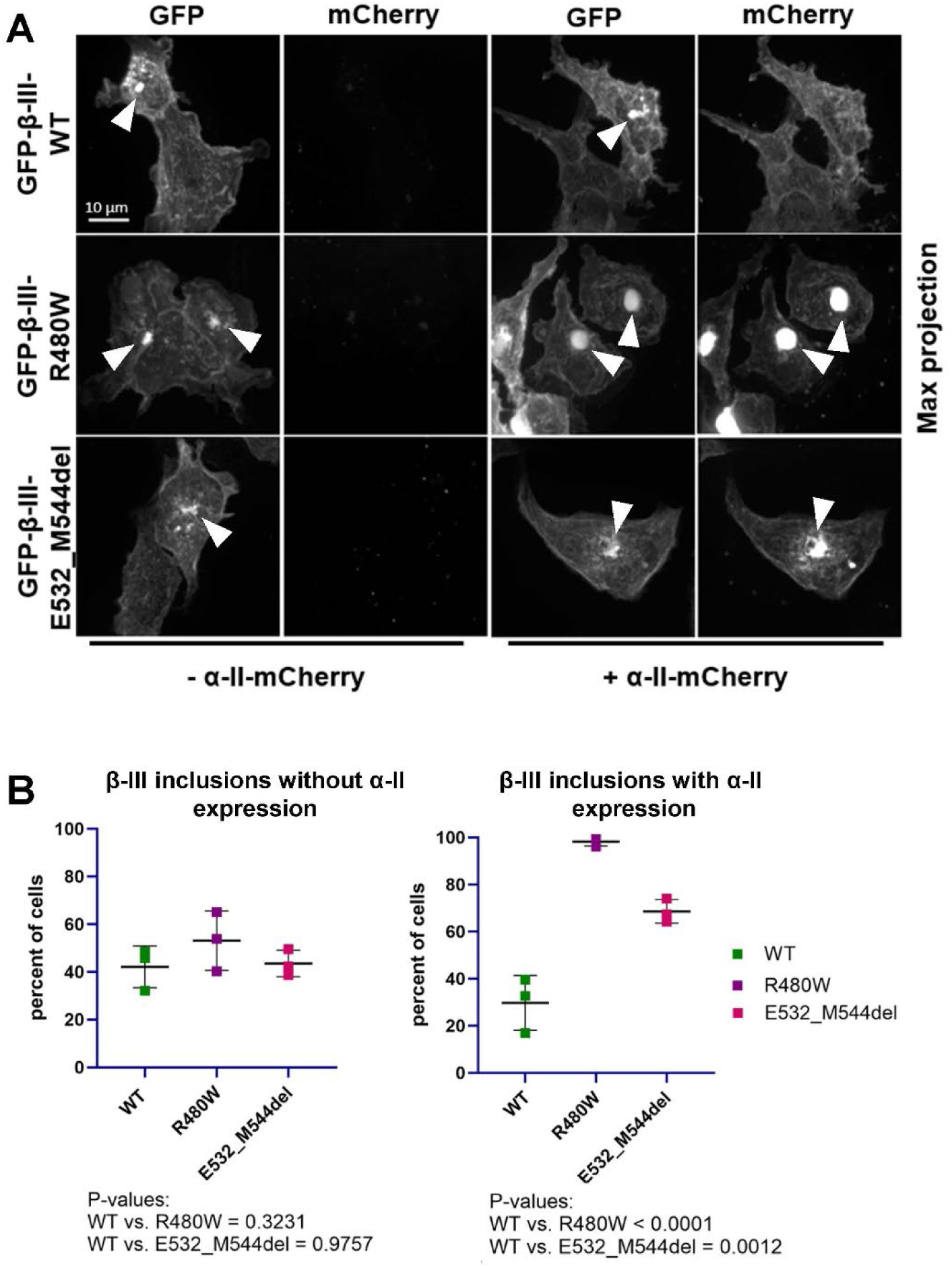
R480W and E532_M544del mutations induce inclusion formation when co-expressed with α-II-spectrin. (A) Representative z-stack max projection images of HEK293T cells showing inclusions containing full-length GFP-β-III-spectrin protein, in the absence (left) or presence (right) of α-II-spectrin-mCherry. In the absence of α-II-spectrin co-expression, wild-type and mutant β-III-spectrin inclusions (white arrows) appear similarly small and irregularly shaped. Co-expression of α-II-spectrin does not change the shape or size of wild-type or E532_M544del β-III-spectrin inclusions. In contrast, co-expression of α-II-spectrin causes the R480W mutant to localize to large, spherical inclusions that contain α-II-spectrin. (B) Quantitation of cells containing spectrin inclusions. In the absence of α-II-spectrin expression, approximately 40-50% of cells show inclusions for wild-type or mutant β-III-spectrin (left). In contrast, with α-II-spectrin co-expression (right), ∼30% of wild-type β-III-spectrin cells show inclusions. The number of cells showing inclusions increases to ∼95% when R480W mutant is co-expressed with α-II-spectrin and to ∼70% when α-II-spectrin is co-expressed with E532_M544del mutant. Statistical significance was determined using Ordinary one-way ANOVA followed by Dunnett’s multiple comparisons test, data represent three independent experiments, with 77-180 cells scored per wild-type or mutant sample, in each replicate. Error bars equal standard deviation.

In contrast, when α-II-spectrin is co-expressed, ∼100% of cells expressing the R480W mutant show inclusions. These inclusions appear larger and more spherical compared to the inclusions observed for R480W in the absence of α-II-spectrin, Fig. 5A. Also, with α-II-spectrin co-expression, for E532_M544del, the percentage of cells showing inclusions (∼70%) was higher than for wild-type (∼40%), Fig. 5B. Like R480W, the E532_M544del inclusions contained α-II-spectrin. However, unlike R480W inclusions, the E532_M544del inclusions appeared small and less regularly shaped (not spherical).

Altogether, these data support that both R480W and E532_M544del proteins physically associate with α-II-spectrin in cells. However, the increased incorporation of both mutants into inclusions, by a mechanism dependent on α-II-spectrin co-expression, supports that both mutations disrupt the α-II/β-III-spectrin complex. In particular, the striking large and spherical inclusions observed for R480W suggest that the mutation grossly alters the α-II/β-III-spectrin complex in a way that is not captured by our FRET, co-IP nor native PAGE binding assays.

### E532_M544del increases β-III-spectrin actin binding

A shared molecular consequence for many ABD-localized SCA5 mutations is increased actin-binding affinity (35,36). Previously, it was shown for β-II-spectrin that attachment of the proximal SRDs to the N-terminal ABD enhances actin binding (34). This indicates that the SRDs contribute to actin binding by a direct or indirect mechanism. Due to proximity of R480W and E532_M544del to the N-terminal ABD, we tested the impact of the R480W and E532_M544del mutations on actin binding. Co-sedimentation assays were performed using a fixed concentration (0.5 µM) of purified β-III-spectrin proteins (amino acids 1-1066) and increasing concentrations of F-actin (0-120 µM). Wild-type β-III-spectrin binds actin with Kd = 24.6 ± 12.0 µM, Fig. 6A. This affinity is higher than the ABD by itself (Kd ∼75 µM) (35,49), supporting a role for the β-III-spectrin SRDs to enhance actin binding. The R480W mutant binds actin with a Kd of 34.9 ± 14.4 µM, indicating that the mutation has no significant impact on β-III-spectrin actin binding. In contrast, over the same actin concentrations, E532_M544del is completely bound to actin. This supports that E532_M544del causes a large increase in actin binding, likely with a sub-micromolar Kd.

**Figure 6.**
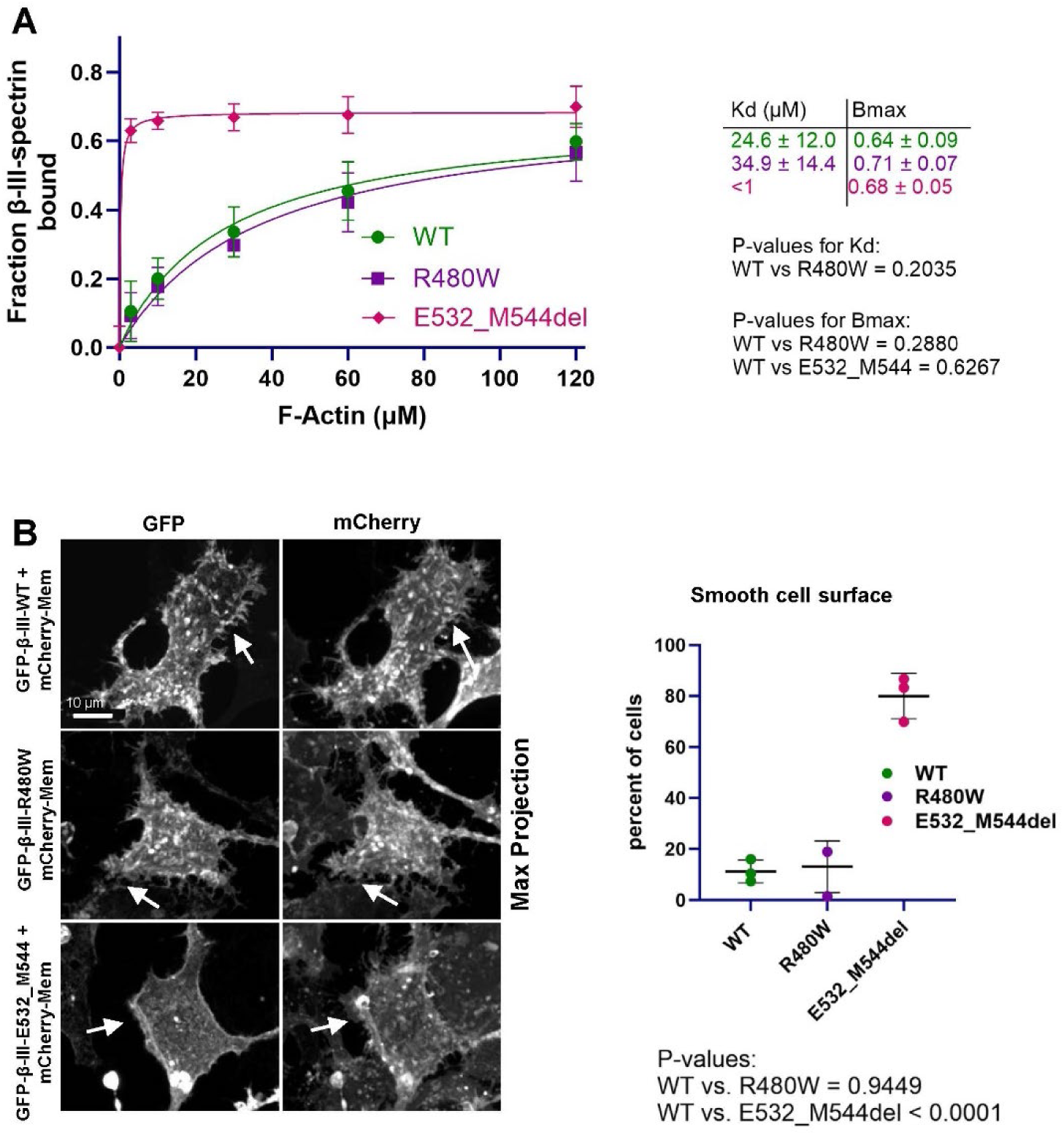
E532_M544del mutation increases β-III-spectrin binding to actin. (A) *In vitro* actin co-sedimentation binding data. Statistical analysis of Kd was performed by unpaired t test. For Bmax, Ordinary one-way ANOVA followed by Dunnett’s multiple comparisons test was used for statistical analysis, n = 4-9 (B) Left, representative z-stack max projection images of HEK293T co-expressing GFP tagged full-length β-III-spectrin proteins and plasma membrane targeted mCherry (mCherry-Mem). Both wild-type and R480W β-III-spectrin proteins localize to filopodia and lamellipodia-like cellular protrusions (white arrow). As a result, images of wild-type and R480W β-III-spectrin proteins give cells a highly textured or “rough” cell surface, similar to images of the mCherry-Mem. The E532_M544del mutant also localizes to the cell cortex. However, the E532_M544del mutant shows reduced localization to filopodia and lamellipodia-like cellular protrusions (labeled with mCherry-Mem). Thus, images of the E532_M544del mutant show a less textured, “smooth” cells surface. Right, quantitation of percentage of cells showing “smooth” cell surface based on β-III-spectrin localization. Approximately 85% of cells expressing the E532_M544del mutant appear smooth compared to ∼20% of cells expressing R480W or wild-type. Statistical significance was determined using Ordinary one-way ANOVA followed by Dunnett’s multiple comparisons test, data represent three independent experiments, with 79-145 cells scored per wild-type or mutant sample, in each replicate. Error bars equal standard deviation.

Previously, we showed that the L253P mutation, which causes high-affinity actin binding (Kd ∼75 nM), causes the full-length β-III-spectrin protein to be absent from lamellipodia and filopodium-like protrusions in HEK293T cells (48). We suggested that increased actin binding limits the pool of free β-III-spectrin available to enter dynamic membrane extensions. To test whether the E532_M544del mutant shows a similar mislocalization to the cellular protrusions, the full-length E532_M544del β-III-spectrin mutant was co-expressed with a mCherry-tagged membrane marker (mCherry-Mem) in HEK293T cells. As reported previously (48), the wild-type β-III-spectrin is detected in filopodium-like and lamellipodia plasma membrane extensions, Fig. 6B. The presence of wild-type β-III-spectrin in these membrane extensions causes cells stained for β-III-spectrin to have a highly textured or “rough” surface. Similar to wild-type, the R480W mutant is present in membrane extensions. In contrast, the E532_M544del mutant is absent from filopodium-like and lamellipodia extensions counter-labelled with mCherry-Mem. The lack of the E532_M544del mutant in cell surface extensions causes cells stained for β-III-spectrin to have a cell surface that appears untextured or “smooth”. Indeed, ∼85% of cells expressing the E532_M544del mutant show a smooth cell surface, compared to ∼10% of cells for wild-type or R480W. These data support that the E532_M544del mutant binds actin with increased affinity in cells, consistent with our co-sedimentation assays showing increased actin binding *in vitro*.

## Discussion

Multiple SCA5 mutations localize to the β-III-spectrin SRDs. However, little is known about the molecular consequences of these mutations. Here we assessed the molecular impact of the SRD2-localized R480W mutation, which is associated with early onset and severe symptoms. We also characterized the SRD3-localized E532_M544del mutation that is associated with late onset and relatively mild symptoms. Based on the position of these mutations in the N-terminal SRDs, we hypothesized that they would impact actin binding and dimerization with α-II-spectrin. Our results support that these mutations differently impact the association of β-III-spectrin with actin and α-II-spectrin. R480W has no impact on actin binding. However, R480W causes the formation of large intracellular inclusions containing the mutant β-III-spectrin and α-II-spectrin. The R480W mutant forms these inclusions only when it is co-expressed with α-II-spectrin, suggesting that R480W grossly alters the structure of α-II/β-III-spectrin complex. In contrast to R480W, the E532_M544del mutation causes β-III-spectrin to bind actin with increased affinity. E532_M544del additionally alters the physical association of β-III-spectrin with α-II-spectrin. Specifically, the E532_M544del mutation causes a loss in coupling of β-III-spectrin SRDs, beginning with SRD3, to the complementary SRDs of α-II-spectrin.

The R480W mutation is located in SRD2, a domain known to be required for nucleating α/β-spectrin dimer through electrostatic interactions (31,32). Our AlphaFold modelling of the α-II/β-III-spectrin nucleation site and the recent cryo-EM analyses of the erythrocyte α-I/β-I-spectrin dimer support that R480 directly contacts α-II-spectrin (44). Thus, we expected that substitution of R480 with a hydrophobic tryptophan residue would alter dimer formation. However, we did not detect a decrease or increase in binding of R480W β-III-spectrin to α-II-spectrin in FRET, co-IP nor native PAGE mobility shift assays, using N-terminal and C-terminal fragments of β-III-spectrin and α-II-spectrin, respectively. Thus, any change in binding of the R480W mutant to α-II-spectrin is below the sensitivity of these assays. Strikingly, co-expression of α-II-spectrin with the R480W β-III-spectrin mutant induces the formation of large, spherical cytoplasmic inclusions containing both the R480W mutant and α-II-spectrin. The dependence of R480W inclusion formation on the co-expression of α-II-spectrin supports that the structure of α-II/β-III-spectrin complex is grossly altered by R480W, even though the mutation did not otherwise cause a detectable change in binding of the two proteins in our other experiments.

Interestingly, we observed that R480W increases β-III-spectrin Tm by ∼1°C compared to wild-type. This increase in stability of the R480W mutant was previously predicted using thermodynamic computational software (15), and experimentally confirmed here. However, the implications of this increased stability are not clear. We speculate, similar to Nuovo et al, that the increased stability may cause a “stiffening” of the β-III-spectrin molecule. The R480W mutant may preferentially populate small number of thermodynamically stable conformations. In contrast, the wild-type protein can sample a greater number of conformational states. It is known that SRDs confer conformational flexibility to spectrin proteins (40,50). A decrease in conformational flexibility might reduce the incorporation (or retention) of the mutant α-II/β-III-spectrin dimer/tetramer into the sub-plasma membrane cytoskeleton, leading to cytoplasmic inclusion formation.

The spherical shape of the R480W inclusions is reminiscent of phase-separated condensates. Interestingly, α-II/β-II-spectrin phase-separated condensates were recently suggested to be present in developing neurites (51). Spectrin condensate formation appeared to be mediated by the spectrin-repeat domains (51). Thus, it is possible that the R480W mutation traps the α-II/β-III-spectrin complex in a biomolecular condensate that is relevant to normal spectrin biology in neurons. Of note, the inclusions caused by R480W appear distinct from the inclusions we previously observed for L253P, which appear as a cluster of small vesicles (48). We suggest that the formation of the R480W inclusions contributes to SCA5 pathogenesis and the associated early onset, severe symptoms. By forming these intracellular accumulations, R480W likely reduces the amount of β-III-spectrin and α-II-spectrin at the plasma membrane, where the wild-type protein normally localizes. In future work, it will be important to study the R480W mutation in an animal model or primary cells to determine if R480W causes inclusion formation at endogenous spectrin protein levels in neurons.

Interestingly, R480 is closely positioned to β-III-spectrin residue R437. These two arginine residues share Y477 as a contact within β-III-spectrin, and both arginine residues contact α-II-spectrin. Several R437 missense mutations have been reported that cause severe symptoms similar to those caused by R480W (16–18). Like R480W, R437Q was predicted to confer structural stability (17). Thus, it will be important to test whether R437 mutations also cause inclusion formation.

Our data show that SRD3-localized E532_M544del impacts β-III-spectrin interaction with both actin and α-II-spectrin. Our co-IP and native PAGE binding experiments with the N-terminal β-III-spectrin fragment (amino acids 1-1066) and complementary C-terminal α-II-spectrin fragment (amino acids 1545-2473) show that the E532_M544del mutant can bind α-II-spectrin. However, our FRET data indicate that the mutation decreases the apposition/coupling of α-II/β-III-spectrin SRDs outside of the high-affinity dimer nucleation site. Our AlphaFold modelling suggests that the E532_M544del introduces a disordered region at the N-terminal half of SRD3 helix A. This disordered region likely acts as a hinge, increasing conformational flexibility of the E532_M544del mutant and interfering with the pairing of downstream α-II/β-III-spectrin SRDs. The E532_M544del mutant can still form a dimer at the nucleation site (SRDs 1-2 of β-III-spectrin), but the C-terminal SRDs do not associate with α-II-spectrin. This unique dimer conformation is supported by our FRET data and the slower migration of the E532_M544del α-II/β-III-spectrin complex in native PAGE compared to wild-type. The uncoupling of SRDs in the spectrin heterodimer would further be expected to interfere with the co-association of two dimers to form a tetramer. A loss in tetramer formation would impede the actin cross-linking function of β-III-spectrin. The E532_M544del mutant inclusions observed in HEK293T cells, when α-II-spectrin is co-expressed, may reflect improperly formed heterodimers/tetramers.

We also found that the E532_M544del mutant causes increased actin binding. The mechanism by which E532_M544del increases actin binding is not clear. It is possible that the E532_M544del causes direct binding of SRDs to actin or that the mutation allosterically enhances actin binding of the N-terminal ABD. In addition, our AlphaFold model showed that the E532_M544del mutation allows the C-terminus to fold back onto the β-III-spectrin N-terminus. It is possible that a novel contact between SRDs and the N-terminal ABD enhances binding of the ABD to actin. Interestingly, a A850G missense mutation in SRD6 of the closely related β-II-spectrin was recently reported to increase actin binding (52).

Numerous ABD-localized mutations, including L253P, increase actin binding (35,36). As we reported for L253P (48), the E532_M544del mutation decreases the localization of β-III-spectrin to filopodium-like and lamellipodia cellular protrusions in HEK293T cells. Further, our prior experiments in *Drosophila* showed that the L253P mutation decreases β-spectrin localization to dendritic extensions (48). Fujishima et al confirmed in cultured Purkinje neurons that the L253P mutation decreases β-III-spectrin dendritic localization (53). Fujishima et al further showed that the E532_M544del mutant was similarly restricted to the soma and absent from Purkinje cell dendrites (53). We suggest that increased actin binding decreases the mobile pool of β-III-spectrin, reducing expansion of the spectrin cytoskeleton into dendrites.

A SCA5 mouse model was previously reported in which a human E532_M544del β-III-spectrin transgene was expressed in Purkinje neurons (54). This SCA5 mouse is ataxic and exhibits thinning of the cerebellar molecular layer containing Purkinje cell dendrites. Interestingly, mGluR1α receptor clustering was found to be reduced in Purkinje cell dendritic spines. This is consistent with a knockout mouse study showing that β-III-spectrin is required for normal spine density and morphology, and the proper localization of other membrane proteins involved in glutamate signalling (55). In the SCA5 mouse model, the E532_M544del mutant β-III-spectrin appears present in dendritic arbores, although quantitation of dendritic localization of the mutant versus wild-type protein was not performed. It is possible that the E532_M544del slows, but does not completely block, the entry of β-III-spectrin into dendrites. We think it is likely that quantitative analysis of the E532_M544del mutant subcellular localization, as performed by Fujishima et al. with cultured neurons (53), will reveal a reduction of dendrite-localized β-III-spectrin in the SCA5 mouse model, at least at early developmental time points. We suggest that either a reduction in E532_M544del mutant localization to dendrites, and/or impaired α-II/β-III-spectrin heterodimer/tetramer formation, is responsible for the decrease in clustering of mGluR1α in dendritic spines.

A second late onset SCA5 mutation that localizes to SRD3 also causes an in-frame deletion (L629_R634delinsW; French kindred). Similar to E532_M544del, the L629_R634delinsW mutant showed reduced localization in the dendrites of cultured Purkinje neurons (53). This similar mislocalization suggests that the L629_R634delinsW mutation also increases actin binding. Moreover, AlphaFold predicts that L629_R634delinsW introduces a disordered region that acts as flexible hinge near the linker connecting SRD3 to SRD4. Thus, we expect that L629_R634delinsW, like E532_M544del, will also disrupt coupling of SRDs in the α-II/β-III-spectrin dimer.

### Experimental procedures Expression constructs

#### GFP and mCherry intermediate constructs

To fuse GFP at the N-terminus of full-length β-III-spectrin, a pcDNA3.1-mEGFP-N intermediate construct was generated as follows: the coding sequence for mEGFP was synthesized by IDT DNA Technologies, digested with KpnI and EcoRI, and inserted into pcDNA3.1(+) vector. To fuse GFP tag at the C-terminus of β-III-spectrin fragment proteins (amino acids 1-1066), pcDNA3.1-mEGFP-C intermediate construct was generated as follows: the coding sequence for mEGFP was PCR amplified using pcDNA3.1-mEGFP (49) as a template, the forward primer CTT GGT ACC ACC ATG GTG AGC and the reverse primer AAA TCT AGA CTA CTT GTA CAG CTC GTC CAT GCC. The reverse primer introduces a stop codon at the 3’ sequence of mEGFP. The PCR product was digested with KpnI and XbaI, and inserted into pcDNA3.1(+) vector.

To fuse mCherry tag at the C-terminus of full-length α-II-spectrin, pcDNA3.1-mCherry-C intermediate construct was generated as follows: the coding sequence for mCherry was PCR amplified using LifeAct-mCherry construct (a gift from Michael Davidson; Addgene plasmid #54491) as a template, the forward primer AAA CTC GAG ATG GTG AGC AAG GGC GAG GAG G and the reverse primer AAA TCT AGA TTA CTT GTA CAG CTC GTC CAT GCC. The reverse primer introduces a stop codon at the 3’ sequence of mCherry. The PCR product was digested with XhoI and XbaI, and ligated into pCDNA3.1(+) vector. To fuse mCherry at the N-terminus of α-II-spectrin fragment protein (amino acids 1545-2473), pcDNA3.1-mCherry-N intermediate construct was generated as follows: the coding sequence for mCherry was PCR amplified using LifeAct-mCherry construct (a gift from Micheal Davidson; Addgene plasmid #54491) as a template, the forward primer AAA GGT ACC ATG GTG AGC AAG GGC GAG GAG G and the reverse primer AAA CTC GAG CTT GTA CAG CTC GTC CAT GCC GCC. The PCR product was digested with KpnI and XhoI, and inserted into pcDNA3.1(+) vector.

#### Bacteria expression constructs

For N-terminal β-III-spectrin-FLAG fragment proteins (amino acids 1-1066), the wild-type and E532_M544del coding sequences were PCR amplified using pcDNA3.1-myc-β-III-spectrin (48) and pUASp-hSPAM (56) as templates, respectively, the forward primer AAA CAC CTG CAA AAA GGT ATG AGC AGC ACG CTG TCA CCC and the reverse primer AAA TCT AGA TTA CTT GTC GTC ATC GTC TTT GTA GTC CCG CCG CGC CTC CCC CAG CGA C. The reverse primer encodes the FLAG epitope to attach a FLAG tag at C-terminus of the β-III-spectrin protein (amino acids 1-1066). The PCR products were digested with AarI and XbaI and were ligated into BsaI digested pE-SUMOpro (LifeSensors). This ligation resulted in pE-SUMO-β-III-spectrin-FLAG wild-type (amino acids 1-1066) or E532_M544del mutant construct. pE-SUMO-β-III-spectrin (amino acids 1-1066) with R480W mutation was generated with site-directed mutagenesis using pE-SUMO-β-III-spectrin-FLAG wild-type (amino acids 1-1066) as a template, the sense primer ACA TCG TGG CCT ACA GCG GCT GGG TGC AGG CAG TGG ACG CC and the anti-sense primer GGC GTC CAC TGC CTG CAC CCA GCC GCT GTA GGC CAC GAT GT.

For C-terminal mCherry-α-II-spectrin-HA protein (amino acids 1545-2473), the 3’ coding sequence of α-II-spectrin was PCR amplified using pcDNA3.1-mCherry-α-II-spectrin (amino acids 1545-2473) (described below for Mammalian expression constructs) as template, the forward primer AAA CAC CTG CAA AAA GGT ATG GTG AGC AAG GGC GAG GAG G and the reverse primer AAA TCT AGA CTA AGC GTA ATC TGG AAC ATC GTA TGG GTA AAC GTT CAC GAA AAG CGA. The forward primer anneals to the 5’ sequence of mCherry, and amplifies the mCherry sequence fused to a coding sequence corresponding to the C-terminal region of α-II-spectrin (amino acids 1545-2473). The reverse primer inserts an HA epitope coding sequence at the 3’ sequence of α-II-spectrin. The PCR product was digested with AarI and XbaI and inserted into pE-SUMOpro vector (LifeSensors) digested with BsaI.

#### Mammalian expression constructs

Full-length pcDNA3.1-mEGFP-β-III-spectrin wild-type and E532_M544del constructs were generated in two steps. First, the 5’ coding sequence (1-2025 bp) for β-III-spectrin wild-type and E532_M544del were PCR amplified using pcDNA3.1-myc-β-III-spectrin template for wild-type (48), pUASp-hSPAM template for E532_M544del (56), the forward primer AAA GAA TTC ATG AGC AGC ACG CTG TCA CCC ACA G and the reverse primer AAA TCT AGA ACC GGT CAG GTC TCG GCC. The PCR products were digested with EcoRI and XbaI restriction enzymes, and inserted into pcDNA3.1-mEGFP-N intermediate construct digested with the same enzymes. This insertion resulted in pcDNA3.1-mEGFP-β-III-spectrin (1-2025 bp) intermediate wild-type and E532_M544del constructs. Second, the 3’ coding sequence (2026-7173 bp) of β-III-spectrin was digested with AgeI and XbaI from the pcDNA3.1-myc-β-III-spectrin construct, and ligated into pcDNA3.1-mEGFP-β-III-spectrin (1-2025 bp) intermediate construct digested with the same restriction enzymes. R480W mutation was introduced using site-directed mutagenesis with the pcDNA3.1-mEGFP-β-III-spectrin (1-2025 bp) intermediate construct as a template, the sense primer ACA TCG TGG CCT ACA GCG GCT GGG TGC AGG CAG TGG ACG CC and the anti-sense primer GGC GTC CAC TGC CTG CAC CCA GCC GCT GTA GGC CAC GAT GT. The PCR resulted in pcDNA3.1-mEGFP-β-III-spectrin (1-2025 bp) intermediate construct with R480W mutation. The β-III-spectrin sequence containing R480W mutation was digested with EcoRI and AgeI from the intermediate construct, and inserted into EcoRI and AgeI digested pcDNA3.1-mEGFP-β-III-spectrin construct generated above.

Full-length pcDNA3.1-α-II-spectrin-mCherry was generated in four steps. First, the SbfI recognition sequence in mCherry was mutated using pcDNA3.1-mCherry-C intermediate construct as a template, the forward primer CAG GAC TCC TCC CTG CAA GAC GGC GAG TTC ATC and the reverse primer GAT GAA CTC GCC GTC TTG CAG GGA GGA GTC CTG. This resulted in pcDNA3.1-mCherry-C construct with mutated SbfI site (pcDNA3.1-mCherry-C-ΔSbfI). Second, the 5’ coding sequence (1-3932 bp) of α-II-spectrin was PCR amplified using pF1KB5523-α-II-spectrin (Kazusa Genome Technologies) as a template, the forward primer AAA GGT ACC ATG GAC CCA AGT GGG GTC AAA GTG C and the reverse primer AAA GAA TTC CCT GCA GGT CTT CTG GTG AC. The PCR products were digested with KpnI and EcoRI, and were inserted into pcDNA3.1-mCherry-C-ΔSbfI construct digested with the same enzymes. This insertion resulted in pcDNA3.1-α-II-spectrin-mCherry-C-ΔSbfI (1-3932 bp). Third, the 3’ sequence (3933-7419 bp) of α-II-spectrin was PCR amplified using pF1KB5523-α-II-spectrin (Kazusa Genome Technologies) as a template, the forward primer AAA CCT GCA GGA AAA GTC CAC AGA GTT AAA C and the reverse primer AAA CTC GAG AAC GTT CAC GAA AAG CGA GCG. The PCR product was digested with SbfI and XhoI, and was inserted into pcDNA3.1-α-II-mCherry-C-ΔSbfI (1-3932 bp) construct digested with the same enzymes. After sequencing the final product (pcDNA3.1-α-II-spectrin-mCherry-ΔSbfI), an insertion mutation was found within mCherry sequence. Thus, mutated mCherry sequence was corrected by swapping in the correct mCherry sequence, obtained from pcDNA3.1-mCherry-C, using XhoI and BsrGI.

For N-terminal β-III-spectrin-GFP proteins (amino acids 1-1066) used for FRET and co-IP experiments, the 5’ sequence of β-III-spectrin wild-type was PCR amplified using the full-length pcDNA3.1-myc-β-III-spectrin (48) as a template, the forward primer AAA GCT AGC ATG AGC AGC ACG CTG TCA CCC and the reverse primer AAA GGT ACC CCG CCG CGC CTC CCC CAG CGA C. The PCR product was digested with NheI and KpnI, and was inserted into pcDNA3.1-mEGFP-C intermediate construct digested with the same enzymes. The final product of this insertion is pcDNA3.1-β-III-spectrin-mEGFP (amino acids 1-1066). For R480W and E532_M544del β-III-spectrin-GFP (amino acids 1-1066) proteins, HindIII and AgeI digestion was performed to isolate β-III-spectrin sequence containing R480W or E532_M544del from pE-SUMO-β-III-spectrin-FLAG constructs generated above. The digestion products were inserted into pcDNA3.1-β-III-spectrin-mEGFP (amino acids 1-1066) digested with the same enzymes. The final products are pcDNA3.1-β-III-spectrin-mEGFP (amino acids 1-1066) R480W and E532_M544del constructs.

For mCherry-α-II-spectrin protein (amino acids 1545-2473), the 3’ coding sequence of α-II-spectrin was PCR amplified using pF1KB5523-α-II-spectrin (Kazusa Genome Technologies) as a template, the forward primer AAA CTC GAG GGA GAA TCT CAA ACC CTC C and the reverse primer AAA TCT AGA TTA AAC GTT CAC GAA AAG CGA G. The PCR products were digested with XhoI and XbaI, and inserted into pcDNA3.1-mCherry-N digested with same enzymes. This insertion resulted in pcDNA3.1-mCherry-α-II-spectrin (amino acids 1545-2473) construct.

PM-GFP was a gift from Tobias Meyer, Stanford University, Stanford, CA (Addgene plasmid #21213). mCherry-Mem was purchased from Addgene (Plasmid #55779). PCR amplifications were performed using iProof-HF polymerase (BioRad). Site-directed mutagenesis was performed using PfuUltra HF polymerase (Agilent). All constructs were sequence verified. The NCBI reference cDNA sequence for β-III-spectrin is NM_006946.4 and α-II-spectrin is NM_001375314.2.

### Protein expression and purification

For N-terminal β-III-spectrin-FLAG (amino acids 1-1066) expression, pE-SUMO-β-III-spectrin-FLAG constructs were transformed into Rosetta2 (DE3) *E. coli* (Sigma). After transformation, single colonies were grown overnight at 250 rpm and 37 °C in a 50 mL LB broth (Miller) supplemented with 100 µg/mL ampicillin and 34 µg/mL chloramphenicol antibiotics. Each overnight culture was expanded into four flasks each containing 250 mL LB broth (Miller) supplemented with the same antibiotics, and grown at 37°C and 250 rpm until the culture OD_550_ reached 0.5-0.7. Protein expression was induced with 0.3 mM IPTG (Fisher bioreagents), and incubated at 16 °C and 250 rpm for 16-24 hours before pelleting.

For purification, the bacteria pellets were resuspended in lysis buffer (50 mM Tris pH 7.5, 300 mM NaCl, 25% sucrose, 1X cOmplete, EDTA-free protease inhibitors (Roche) and 4:1 (w/v) Lysozyme (GoldBio)), and stirred at 4 °C for 30 minutes. Further protein extraction was achieved with sonication at 15 seconds on and 45 seconds off cycles for total time of 3 minutes at 45% amplitude, using a Branson 250 sonifier with tip size of ½ inch. Lysate supernatants were collected after centrifugation at 40,905 x g and 4°C using Sorvall SS-34 rotor for 30 minutes. The supernatants were clarified using 0.45 µm disk filters, and loaded into Poly-Prep chromatography column (BioRad) packed with equilibrated Ni-NTA agarose beads (Invitrogen). Following three washes with 50 mM Tris pH 7.5, 300 mM NaCl and 20 mM imidazole buffer, SUMO-β-III-spectrin-FLAG proteins were eluted off the columns with 50 mM Tris pH 7.5, 300 mM NaCl and 150 mM imidazole buffer. After adding a 1:1 volume ratio of 25 mM Tris pH 7.5, 150 mM NaCl, and 5 mM 2-Mercaptoethanol buffer to the eluted proteins, SUMO tag was digested with 1:100 mass ratio of Ulp1 SUMO protease: total eluted proteins for 30 minutes at 4°C. Then, digested proteins were loaded into 10,000 MWCO Slide-A-Lyzer cassettes (Thermo Scientific) for buffer exchange into 50 mM Tris pH 7.5 and 150 mM NaCl, and incubated at 4 °C stirring for 4-6 hours to remove 2-mercaptoethanol. After dialysis, the proteins were loaded into Poly-Prep columns packed with equilibrated anti-FLAG M2 affinity gel beads (Sigma) at 4 °C. After several washes with 50 mM Tris pH 7.5 and 150 mM NaCl buffer, β-III-spectrin-FLAG (amino acids 1-1066) proteins were eluted with 10 mM Tris pH 7.5, 150 mM NaCl and 200 ng/µL FLAG peptide (Sigma) buffer. Final buffer exchange into 10 mM Tris pH 7.5, 150 mM NaCl, 2 mM MgCl_2_ and 1 mM DTT buffer was performed using 10,000 MWCO Slide-A-Lyzer cassettes stirring overnight at 4 °C. Pure β-III-spectrin-FLAG proteins were supplemented with 25% sucrose before snap-freezing and storage at −80 °C.

A similar expression protocol was followed for C-terminal mCherry-αII-spectrin-HA fragment (amino acids 1545-2473). For protein purification, SUMO-mCherry-α-II-spectrin-HA protein was purified by Ni-NTA beads, followed by cleavage of the SUMO tag using Ulp1 protease. The digest was then loaded onto a GE S100 size exclusion column equilibrated in 10 mM HEPES pH 7.5, 150 mM NaCl, 2 mM MgCl_2_ and 1 mM DTT buffer, and fractions containing mCherry-α-II-spectrin-HA collected.

### Circular dichroism spectroscopy

Purified β-III-spectrin-FLAG fragment proteins (amino acids 1-1066) were thawed and clarified at 100,000 x g and 4 °C for 15 minutes using TLA 100.3 rotor (Beckman). Bradford assay was performed to determine protein concentration. β-III-spectrin-FLAG proteins (amino acids 1-1066) were diluted to a final concentration of 150 ng/mL in 10 mM Tris pH 7.5, 150 mM NaCl, 2 mM MgCl_2_, 1 mM DTT buffer. Jasco J-810 spectropolarimeter with Peltier temperature controller was used to collect absorption profiles and thermal denaturation data as was described previously (38). Absorption data were reported as mean residue ellipticity (degrees cm^2^ dmol^-1^) after converting the raw data (38). Thermal denaturation data were measured at 222 nm, and melting temperatures were calculated as described previously using Prism 10 (GraphPad) and non-linear regression to fit a two-state transition equation (38).

### Live-cell FRET assay

HEK293T cells (ATTC) were either transfected with 0.5 µg pcDNA3.1-β-III-spectrin-mEGFP (amino acids 1-1066) FRET donor constructs (D) only, or co-transfected with 2 µg pcDNA3.1-mCherry-α-II-spectrin (amino acids 1545-2473) FRET acceptor construct (A) following Lipofectamine 3000 protocol (ThermoFisher Scientific) in a 6-well plate. Twenty-four hours following transfection, cells were harvested and prepared for GFP fluorescence lifetime measurement as previously described (49). FS5 spectrofluorometer equipped with an EPL-472 pulse laser with 5 MHz frequency (Edinburgh Instruments) was used to collect and calculate GFP lifetime (τ) in donor (D) and donor acceptor (DA) samples. FRET efficiency was calculated using the following equation:

### Co-immunoprecipitation assay

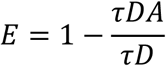

For β-III-spectrin-GFP (amino acids 1-1066) and mCherry-α-II-spectrin (amino acids 1545-2473) co-immunoprecipitation (co-IP) assay, 0.5 µg of pcDNA3.1-β-III-spectrin-mEGFP constructs were co-transfected with 2 µg pcDNA3.1-mCherry-α-II-spectrin in HEK293T cells in a 6-well plate, using Lipofectamine 3000, as described for FRET assays. Twenty-four hours post transfection, cells were washed once with 1X DPBS (Gibco), resuspended in ice-cold lysis buffer (10 mM Tris pH 7.5, 150 mM NaCl, 320 mM sucrose, 2 mM EDTA, 1% Triton X-100, 0.5% IGEPAL CA-630, 0.1% SDS, 1X cOmplete mini, EDTA-free protease inhibitor (Roche)) and incubated on ice for 30 minutes with gentle swirling every 10 minutes. Supernatants were collected after spinning the lysates at 16,000 x g and 4 °C for 10 minutes. A small volume of clarified supernatants was mixed with 4x Laemmli sample buffer (BioRad) and saved at −20 °C for lysate/input analysis on western blots.

To precipitate β-III-spectrin-GFP proteins, GFP-Trap magnetic agarose (Chromotek) beads were first equilibrated with 10 mM Tris pH 7.5, 150 mM NaCl, 2 mM EDTA buffer, and then were incubated with the supernatant samples rotating at 4 °C for 1 hour. After several washes with 10 mM Tris pH 7.5, 150 mM NaCl, 2 mM EDTA buffer, IP samples were eluted into 4x Laemmli sample buffer and heated to 95 °C for 5 minutes.

For western blot analysis, IP and lysate/input samples were resolved by SDS-PAGE and transferred to Immobilon-FL membranes (Millipore). Membranes were blocked with 1X PBS-L (48) containing 1% casein (Hammarsten; Affymetrix). Primary antibodies were diluted in 1X PBS-L, 1% casein and 0.1% Tween 20 buffer, and incubated with the membranes overnight at 4 °C. Primary antibodies included: anti-GFP (1:2000; Millipore), anti-α-II-spectrin C-11 (1:200; Santa Cruz); anti-mCherry (1:2000; Invitrogen). Membranes were washed with 1X PBS-L containing 0.1% Tween 20. Secondary antibodies were diluted to 1:5000 in 1X PBS-L supplemented with 1% Casein, 0.1% Tween 20 and 0.01% SDS, and incubated with the membranes for 1 hour at room temperature. Secondary antibodies used include anti-mouse IRDye 800CW (LI-COR Biosciences) and anti-rabbit IRDye 680RD (LI-COR Biosciences). Membranes were washed with 1X PBS-L containing 0.1% Tween 20 and imaged on a Sapphire scanner (Azure) using 680 nm and 800 nm channels. Fluorescence protein band intensities were quantified using ImageStudio Lite (LI-COR Bioscience) as described previously (49).

### Native PAGE mobility shift assay

The concentration of purified β-III-spectrin-FLAG fragments (amino acids 1-1066) and mCherry-α-II-spectrin-HA fragment (amino acids 1545-2473) proteins was determined by Bradford assay. The proteins were diluted to 0.5 µM in a buffer containing 10 mM HEPES pH 7.5, 150 mM NaCl, 2 mM MgCl_2_, 1 mM DTT. β-III-spectrin-FLAG proteins and mCherry-α-II-spectrin-HA were either incubated separately or combined for 1 hour at room temperature. Then, samples were mixed with 1.5x Native Sample Buffer (Bio-Rad). The samples were loaded onto an 8% acrylamide gel. Native PAGE electrophoresis was performed in ice-cold 1X Tris-glycine (BioRad) running buffer at constant voltage of 75 V for 1 hour and then constant current of 0.01 Amp for 3.5 hours. Following staining of gels with Coomassie R-250, gels were imaged using a Sapphire scanner (Azure). Fluorescence protein band intensities were quantified using ImageStudio Lite (LI-COR Bioscience) as described previously (49).

### Fluorescence subcellular localization assays

Glass coverslips (22 x 22 mm; #1.5; Thermo Fisher Scientific) were cleaned with 70% ethanol, placed in 6-well plate and washed with Ultrapure water (Gibco). The coverslips were incubated with 0.1 mg/mL poly-D-lysine, 70–150 kDa (Sigma) solution for 20 minutes at room temperature. Poly-D-lysine solution was discarded, coverslips washed multiple times with Ultrapure water and sterilized under UV light for 1 hour. HEK293T cells were plated on the coated glass coverslips at 0.3x 10^6^ cells per well, and incubated for 2-4 hours at 37°C and 5% CO2. The cells were co-transfected using Lipofectamine 3000 with 0.5 µg full-length pcDNA3.1-mEGFP-β-III-spectrin or 0.25 µg PM-GFP with or without 0.5 µg full-length pcDNA3.1-α-II-spectrin-mCherry, or with 0.25 µg mCherry-Mem. Twenty-four hours post transfection, the cells were washed with 1X DPBS and fixed with 4% paraformaldehyde in 1X DPBS for 15-30 minutes at room temperature. The cells were washed again with 1X DPBS, and mounted on a 1 mm microscope slide (VWR) using Prolong Diamond antifade with DAPI (Invitrogen). After 24 hours, the cells were imaged using a HC PL APO 63x/1.40-0.60 OIL oil-immersion objective on a Leica DMi8 inverted microscope equipped with X-Light V2 spinning-disk (CrestOptics), LDI laser unit (89 North) and Photometrics Prime 95B CMOS camera. Z-stack images were processed using ImageJ (NIH) or Photoshop software (Adobe). Representative intensity vs distance profiles were obtained using ImageJ line tool and RGB Profile plugin. Pearson correlation coefficient was calculated in Prism 10 (GraphPad) after measuring GFP and mCherry intensities using the Line and Plot Profile tools in ImageJ. Images of cells expressing GFP-β-III-spectrin proteins were visually scored for inclusions or smooth cell surface.

### Actin co-sedimentation assay

Purified β-III-spectrin-FLAG proteins (amino acids 1-1066) were thawed and clarified at 43,000 rpm using TLA-100.3 rotor (Beckman). Actin was extracted from rabbit muscle acetone powder (Pal-Freez Biologicals) and polymerized into F-actin as described previously (49). Bradford assay was performed to measure the concentration of the proteins. β-III-spectrin-FLAG proteins were diluted to 0.5 µM and mixed with varying concentrations of F-actin (3-120 µM) in F-buffer (10 mM Tris pH 7.5, 150 mM NaCl, 0.5 mM ATP, 2 mM MgCl2, 1 mM DTT). For 2-point standard curve, β-III-spectrin-FLAG proteins were diluted to 0.5 µM and 0.175 µM. The samples were incubated at room temperature for 30 minutes. Then, the samples were centrifuged at 50,000 rpm and 25 °C for 30 minutes using TLA-100 rotor (Beckman) to pellet F-actin and bound β-III-spectrin-FLAG proteins. Supernatant samples containing unbound β-III-spectrin-FLAG proteins were collected and mixed with 4x Laemmli sample buffer. Supernatant samples were separated by SDS-PAGE using 12% acrylamide gels. Gels were stained with Coomassie R-250, and imaged using a Sapphire fluorescence scanner (Azure). Fluorescence protein band intensities were quantified using ImageStudio Lite software (LICOR), and dissociation constant (Kd) and Bmax calculated in Prism 10 (GraphPad) using the equation: Y = Bmax*X/(Kd + X), where Y is the bound protein fraction and X is the free F-actin fraction (57).

## Data availability

All data are contained within the manuscript.

## Acknowledgments

We thank Anjelika Gasilina at NIH and Emma Cismas at Oakland University for reviewing the manuscript.

## Author contributions

Adam W. Avery conceived and designed experiments. S.A.D., N.R.B., Abeer W. Abukamil, A.N.A. A.R.K., A.E.A., Adam W. Avery. acquired and interpreted data. Adam W. Avery and S.A.D prepared the manuscript.

## Funding and additional information

This work was supported by the NIH grant R15NS116511 to Adam W. Avery.

